## Supplementary Information for "Genomic variation of a keystone forest tree species reveals signals of local adaptation despite high levels of phenotypic plasticity"

**This file includes:**

Supplementary Figures 1-24

Supplementary Table 4

Legends for Supplementary Tables 1-3 and 5-9

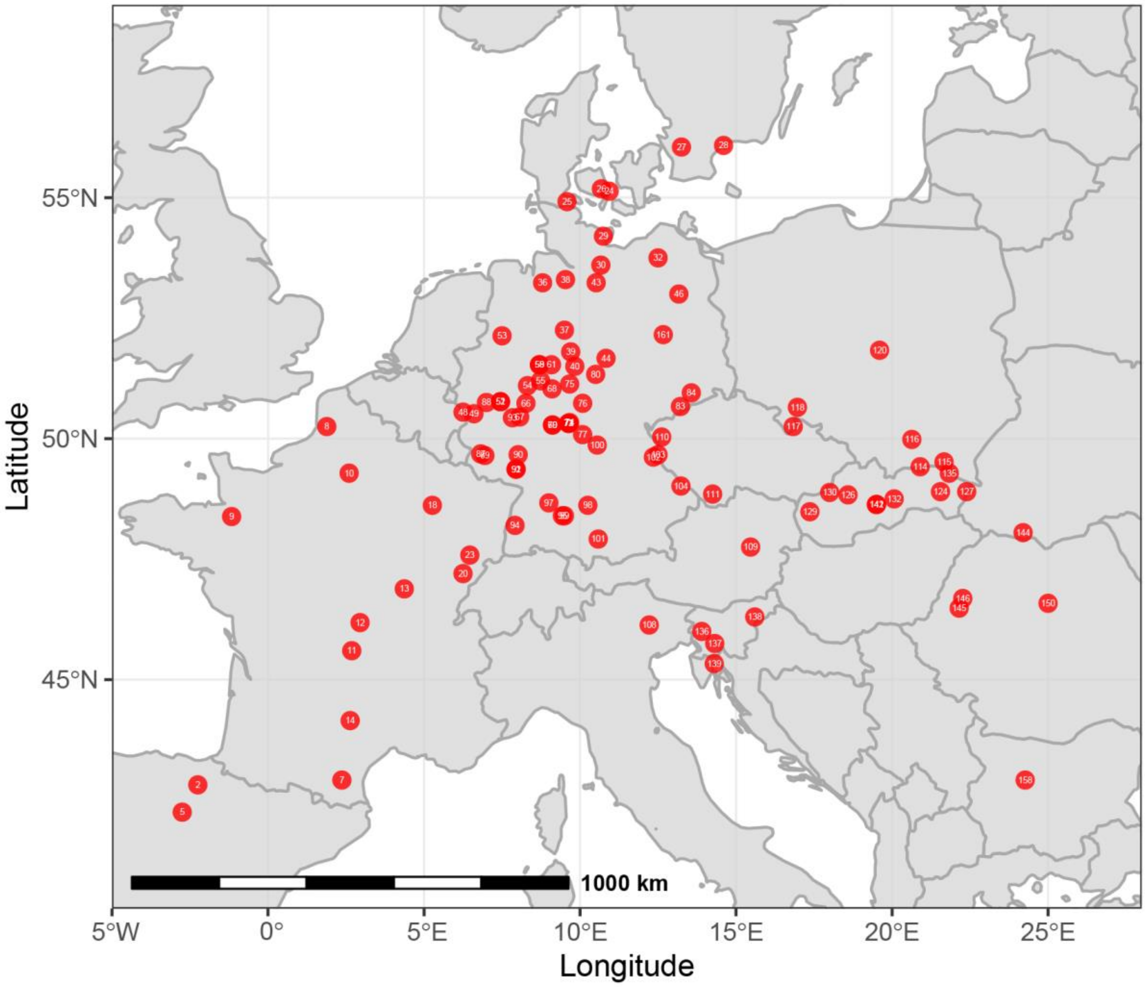


**Supplementary Figure 1**

**Populations (provenances) used for the genomic analyses in this study**

Red circles indicate geographic origins of the 100 populations used in this study (Supplementary Table 1). These populations are part of an international provenance trial, consisting of 38 common gardens distributed across Europe[54], 23 of which were planted in 1995. White numbers indicate provenance codes.

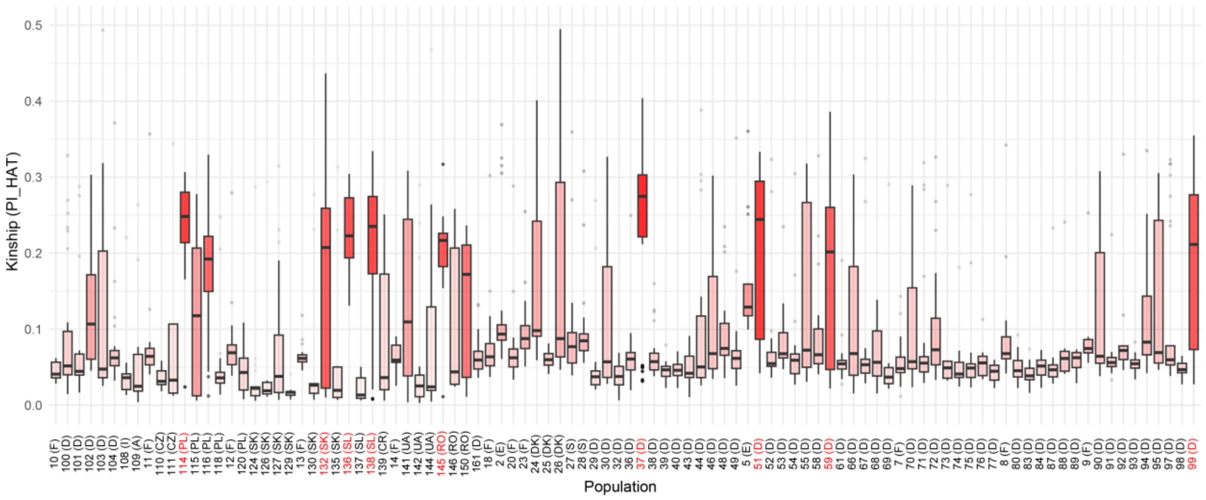


**Supplementary Figure 2**

**The individuals sampled for each population exhibit different levels of kinship**

Boxplots show all pair-wise relationships, that is PI_HAT values calculated with plink as the proportion of identity by descent (IBD) specifically P(IBD=2) + 0.5*P(IBD=1), between individuals of one population[62]. Populations with an average above 0.2 are indicated by red letters. Provenance IDs and countries of origin are shown on the x-axis. Some populations exhibit high levels of pair-wise relationships indicating a limited number of individuals involved in the seed harvest of those populations.

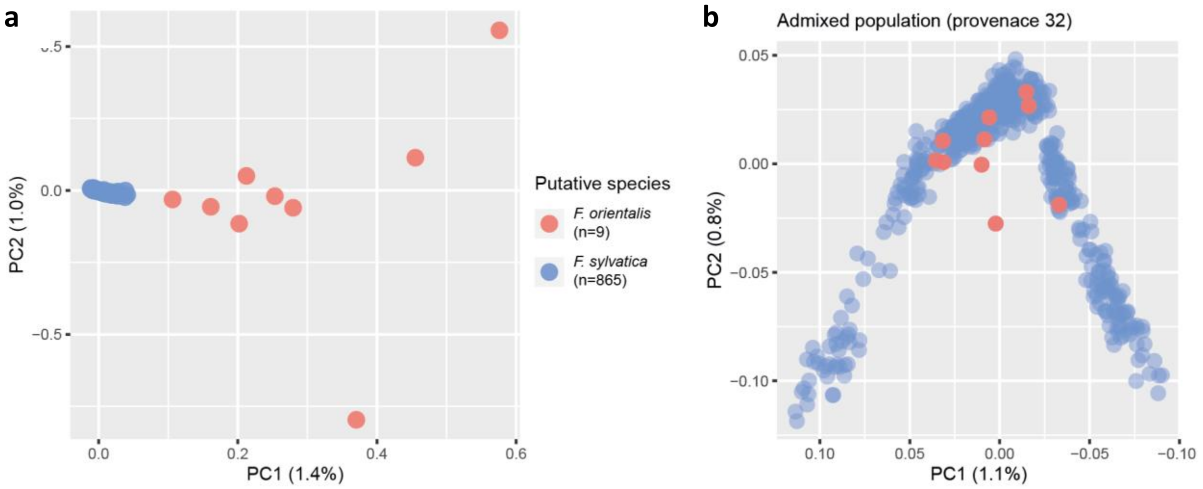


**Supplementary Figure 3**

**One divergent and one admixed population were excluded**

(**a**) The population from Bulgaria (provenance 158) (red) appears highly divergent from all other individuals (blue) and may not represent *Fagus sylvatica* but a hybrid with the sister species *F. orientalis* instead. (**b**) One population from Northern Germany (provenance 32) (red, n=10) appears admixed and may thus represent a seed orchard with material from different parts of the distribution range. None of the other 98 populations (blue, n=855) showed such a pattern.

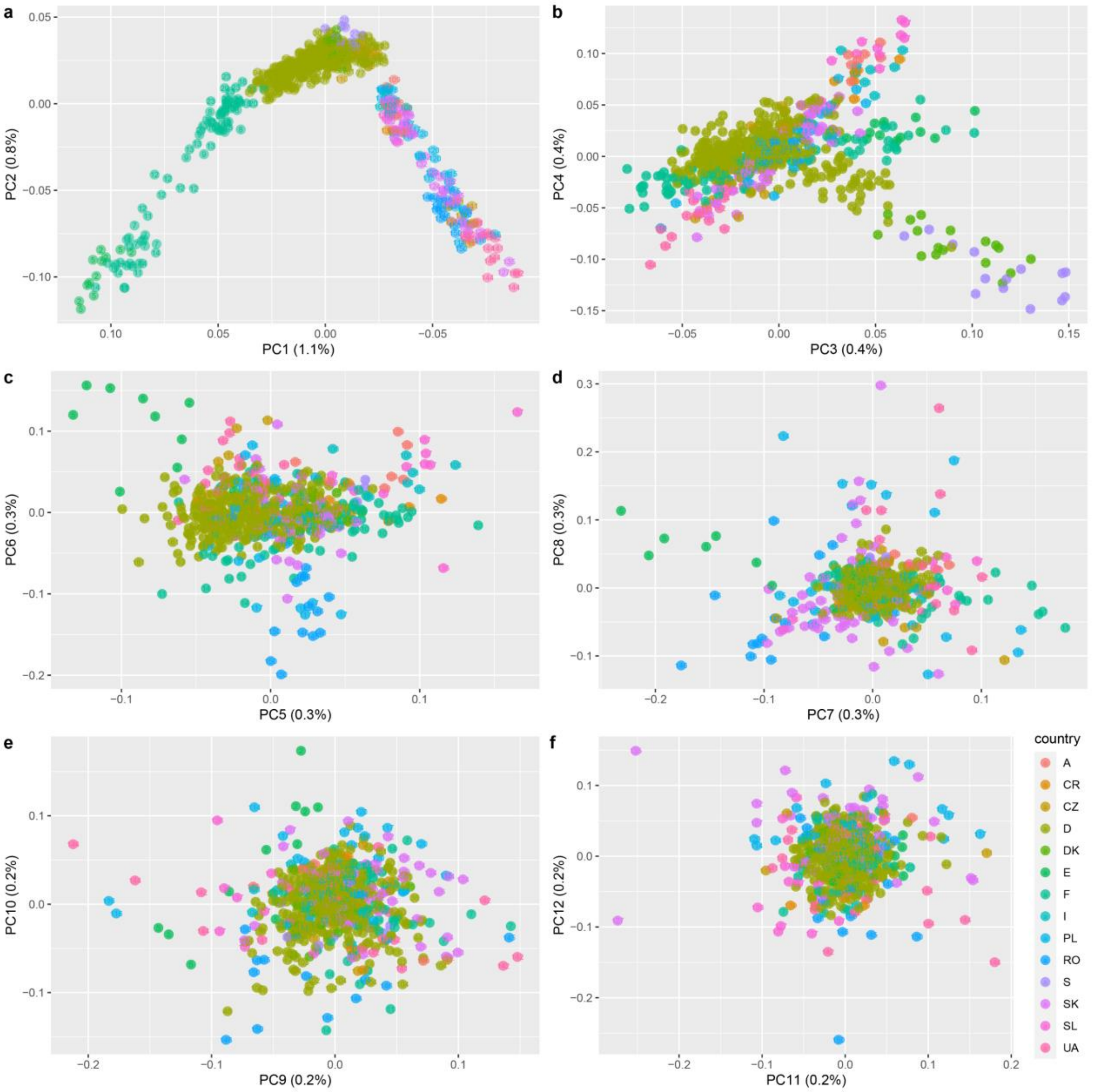


**Supplementary Figure 4**

**Principle component analysis (PCA) shows marked geographic structure up to PC6**

(**a-f**) PCA of 540k independent (LD<0.2) genome-wide variants in 653 largely unrelated (less than 2$^{nd}$ degree) individuals from 98 populations colored by countries of origin shows correlation between genetics and geography for PC1 to PC6, despite small values of variance explained (1.1-0.3%).

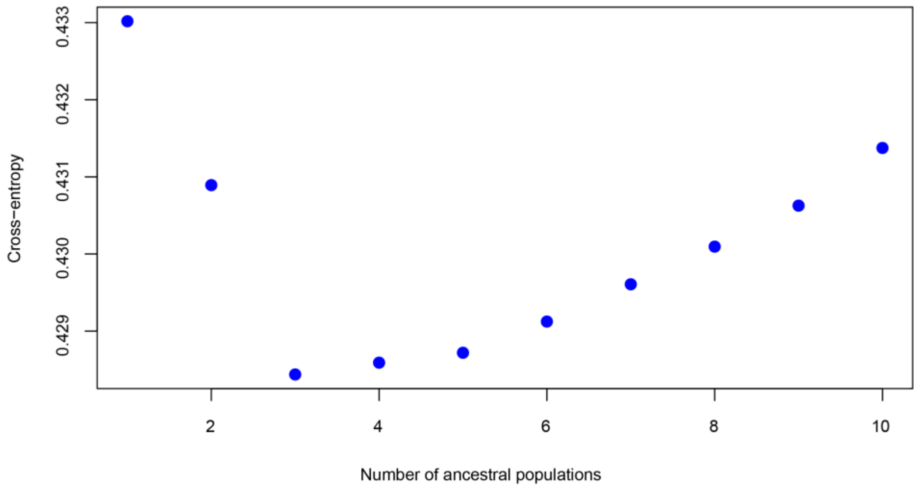


**Supplementary Figure 5**

**Cross-entropy of ancestry coefficients analysis indicates three major ancestral genetic clusters**

The sparse non-negative matrix factorization (snmf) method, which we ran for different numbers of clusters (K=1 to 10) using the 540k independent genome-wide variants in the 653 largely unrelated (less than 2nd degree) individuals from the 98 populations exhibited the lowest cross-entropy for K=3 indicating the presence of three main ancestral genetic clusters[30].

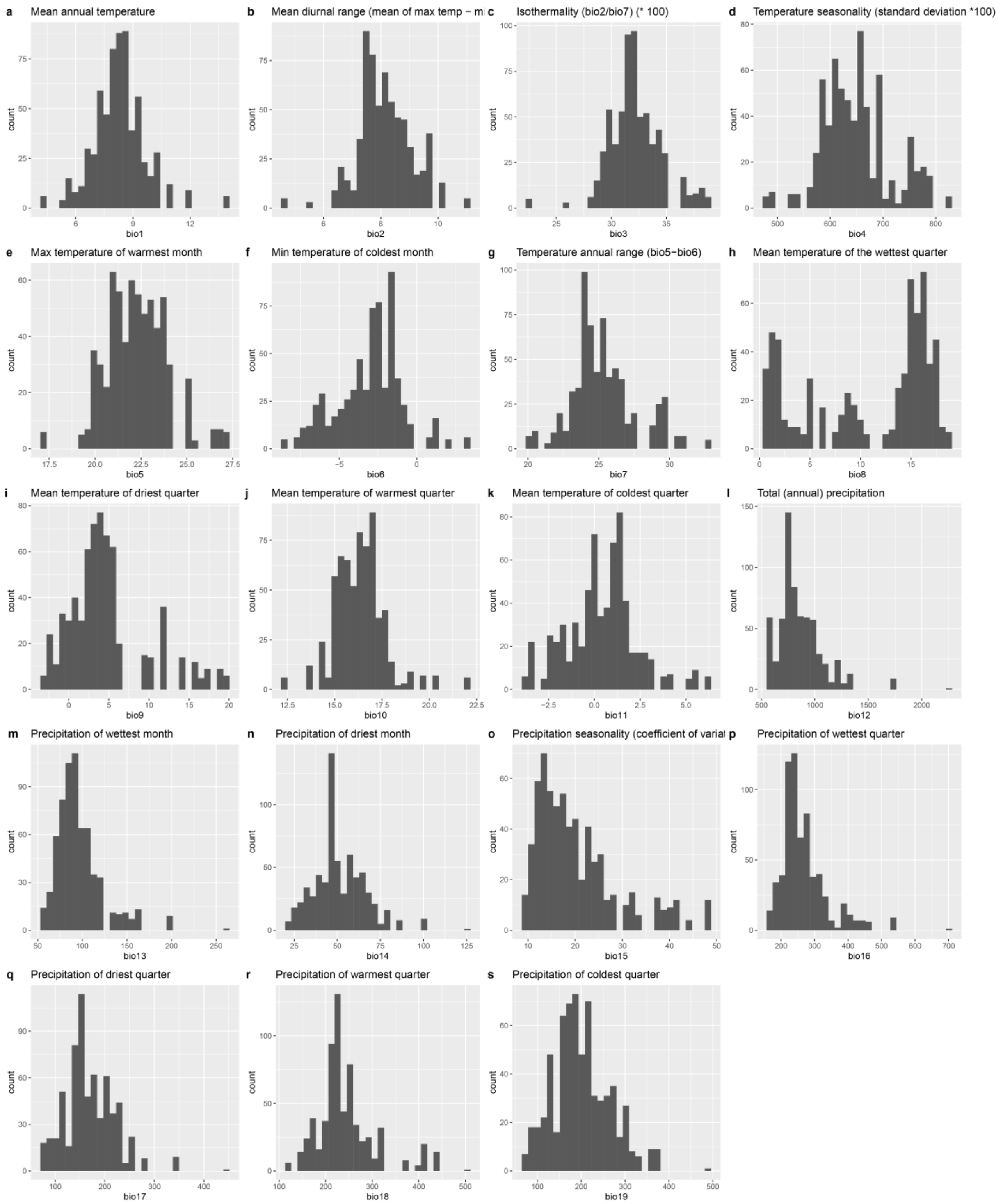


**Supplementary Figure 6**

**Bioclimatic variables exhibit broad and mostly normal distribution across the 98 populations analyzed**

(**a-s**) Histograms show the distribution of the 19 bioclimatic variables from the WorldClim v2.1 database[31] with a 5x5 km resolution across the 98 beech populations analyzed.

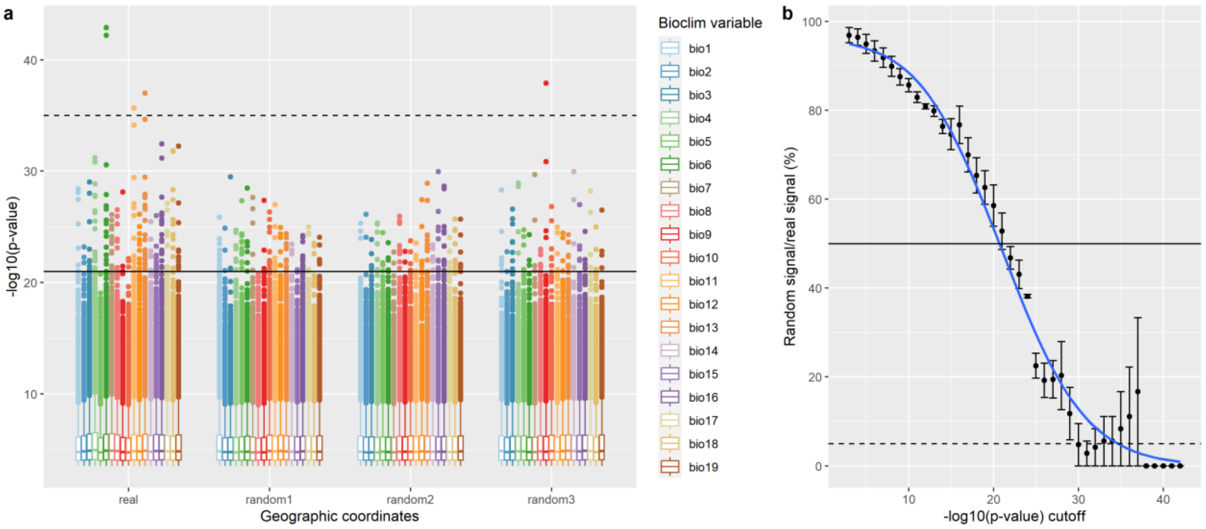


**Supplementary Figure 7**

**Data randomization demonstrates statistically significant but non-causal signal in genotype-environment association analysis using LFMM**

(**a**) We ran the latent factor mixed model (lfmm) analysis[33] using not only the real data but also three random datasets, where the population coordinates and thus the bioclimatic variables were randomly assigned. Boxplot show *p*-values for each of the 19 bioclim variables for each of the four runs (real and random1-3), excluding variants with FDR-corrected *p*-values>0.01. Two variants, especially associated with variation in bio6 but also bio11 and bio13 (also see Supplementary Fig. 8) in the real data, stood out from the rest. Dashed and solid horizontal lines indicate significance thresholds for which the likelihood of being exceeded with random data compared to the real data is 5% (number of associations with random data/number of associations with real data = 1/20) or 50%, respectively, as determined by comparison of the *p*-value distributions between real and random data shown in (b). (**b**) Significance thresholds were defined by assessing the ratio of random vs. real signal (i.e., the number of associations identified with random data divided by the number of associations identified with the real data) in bins of increasing size, starting with the most significant associations and then adding less significant ones consecutively in -log10(*p*-value)-steps of one (-log10(*p*) bins = 42-43, 41-43, 40-43, 39-43 etc.). The blue line shows a polynomial regression of the means, the error bars the standard error between the three random runs.

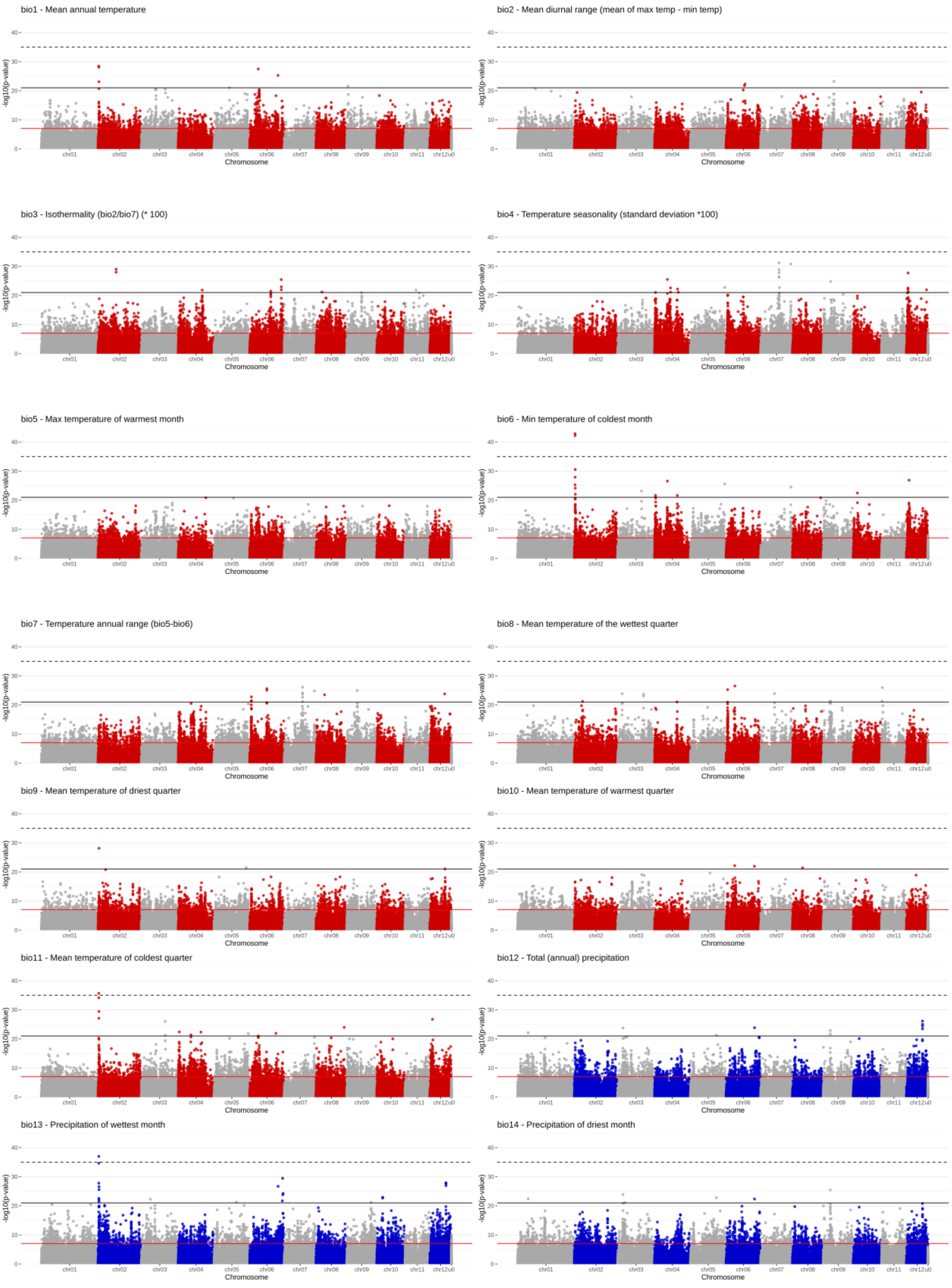
bio1 - Mean annual temperature
bio2 - Mean diurnal range (mean of max temp - min temp)
bio3 - Isothermality (bio2/bio7) (* 100)
bio4 - Temperature seasonality (standard deviation *100)
bio5 - Max temperature of warmest month
bio6 - Min temperature of coldest month
bio7 - Temperature annual range (bio5-bio6)
bio8 - Mean temperature of the wettest quarter
bio9 - Mean temperature of driest quarter
bio10 - Mean temperature of warmest quarter
bio11 - Mean temperature of coldest quarter
bio12 - Total (annual) precipitation
bio13 - Precipitation of wettest month
bio14 - Precipitation of driest month
-log10(p-value)
0
10
20
30
40
chr01
chr02
chr03
chr04
chr05
chr06
chr07
chr08
chr09
chr10
chr11
chr12u0
Chromosome

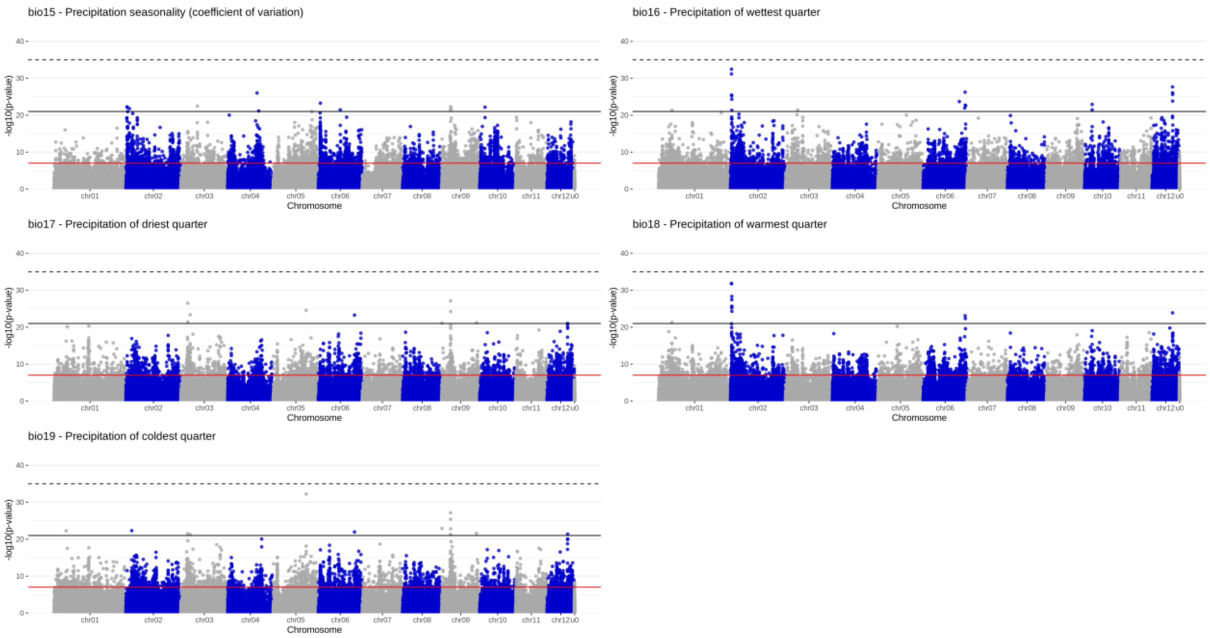


**Supplementary Figure 8**

**Genotype-environment associations for all 19 bioclimatic variables using LFMM.**

Manhattan plots of 540k independent genome-wide variants in 653 individuals from 98 populations across the distribution range. The significance of the associations of each variant along the 12 beech chromosomes and the unplaced contigs (indicated as u0) is shown by the -log10 *p*-values from the LFMM analyses. Dashed and solid black horizontal lines indicate significance thresholds for which the likelihood of being exceeded with random data compared to the real data is 5% (number of associations with random data/number of associations with real data = 1/20) or 50%, respectively, as determined by comparison of the p-value distributions between real and random data as shown in Supplementary Fig. 7. Red horizontal line shows 5% Bonferroni significance threshold. The SNPs of every other chromosome are colored in red for temperature-related bioclimatic variables and in blue for precipitation-related variables.

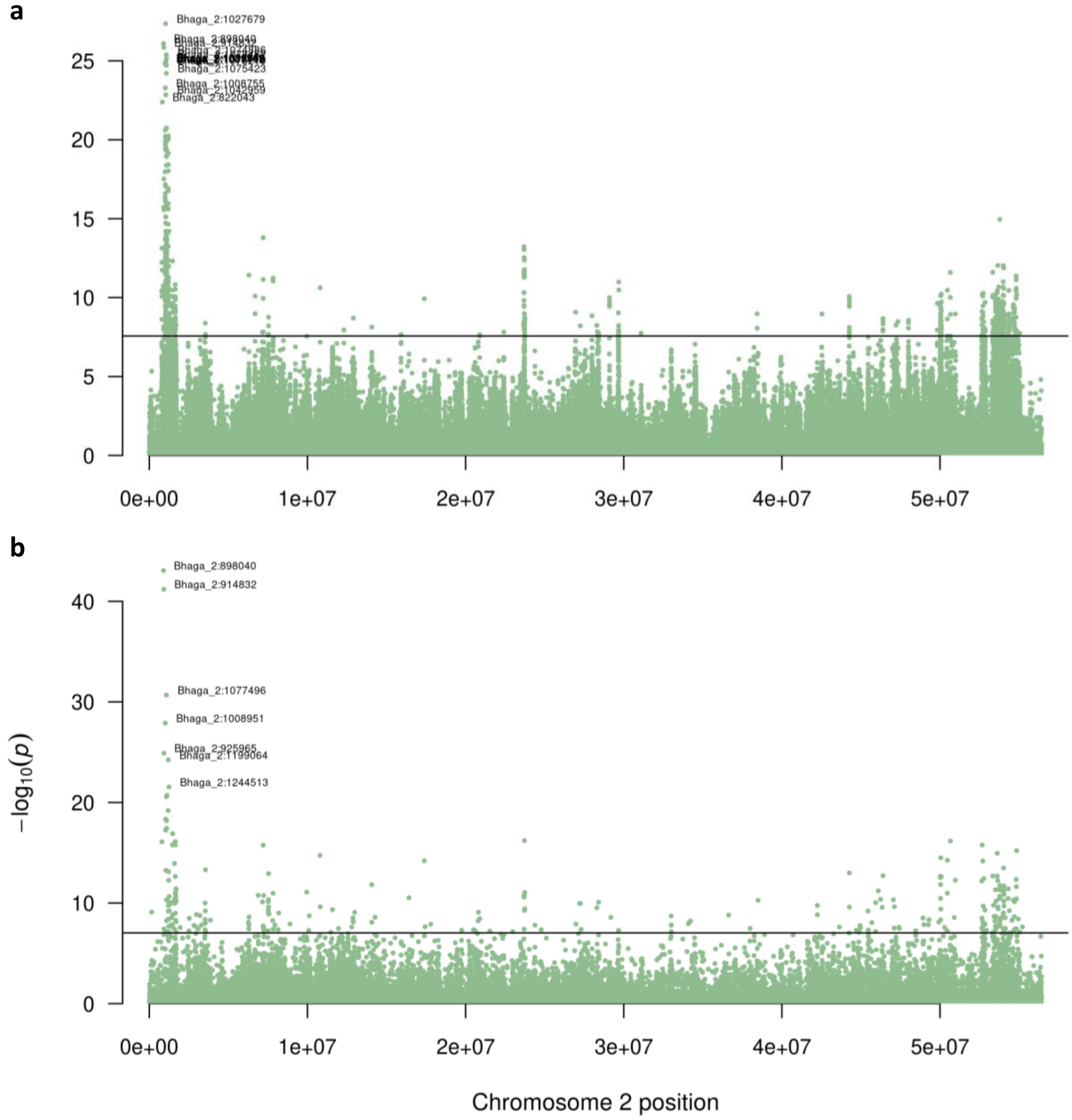


**Supplementary Figure 9**

**Genotype-environment association for winter cold using all variants or only LD-pruned variants.**
(**a**,**b**) Manhattan plots show the significance of variants associated with winter cold in LFMM analyses along chromosome 2 by the -log10 *p*-values using all 3.68 million (a) or only the 540k LD-pruned (b) variants in 653 individuals from 98 populations. Horizontal lines indicate Bonferroni significance thresholds. Variant IDs (Chromosome:position) are depicted for the most significant variants.

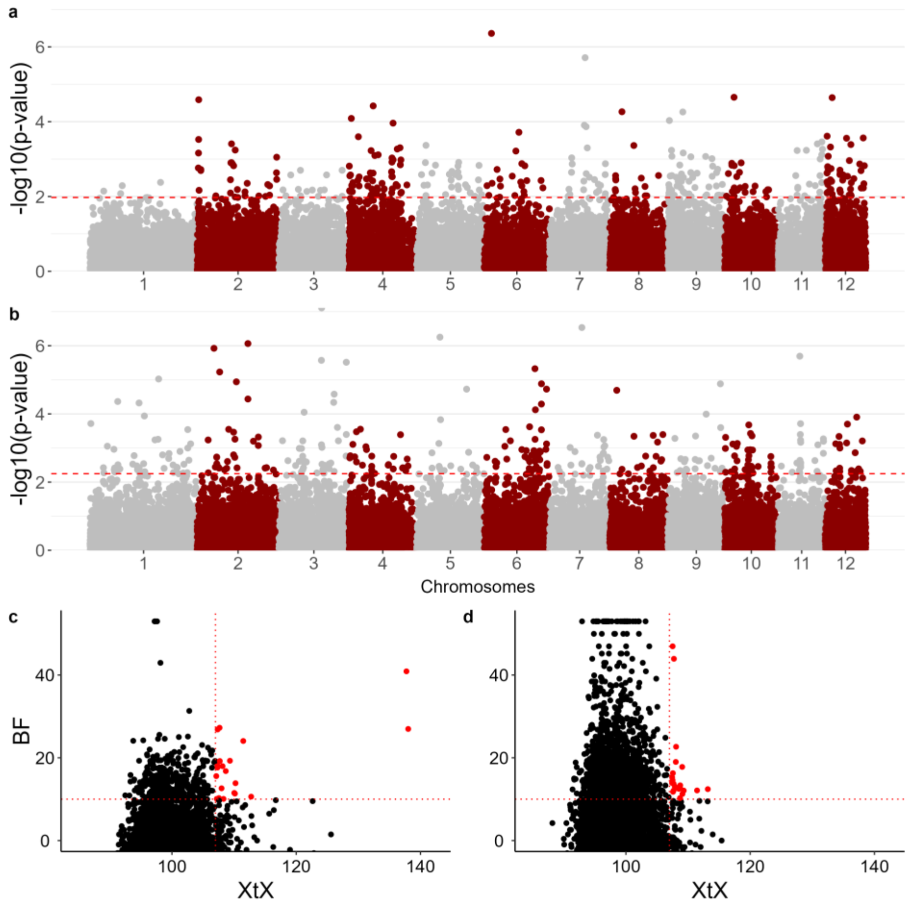


**Supplementary Figure 10**

**Data randomization demonstrates statistically significant but non-causal signal in GEA analysis using WZA and BayPass**

(**a**,**b**) Manhattan plots of 10 kb genome-wide windows in 653 individuals from 98 populations. Chromosomes are distinguished by alternating colors for enhanced visualization. Odd-numbered chromosomes are represented in gray, while even-numbered ones are depicted in red. The significance of the association of each variant along the 12 chromosomes with the minimum temperature of the coldest month (bio6) of the population origins is indicated by the -log10 *p*-values of the WZA scores on the y-axis. The red dashed horizontal lines indicate 99th percentile thresholds for real (a) and randomized (b) data. (**c**,**d**) The XtX genetic differentiation value as a function of the Bayes Factor (BF) of the association with the same bioclimatic variable as in (a) and (b). The red dashed horizontal lines indicate a 10 BF threshold, while vertical lines represent a 107 XtX threshold based on the pseudo-observed data (POD). Red dots represent the outliers considering XtX and BF thresholds for real (c) and randomized data (d).

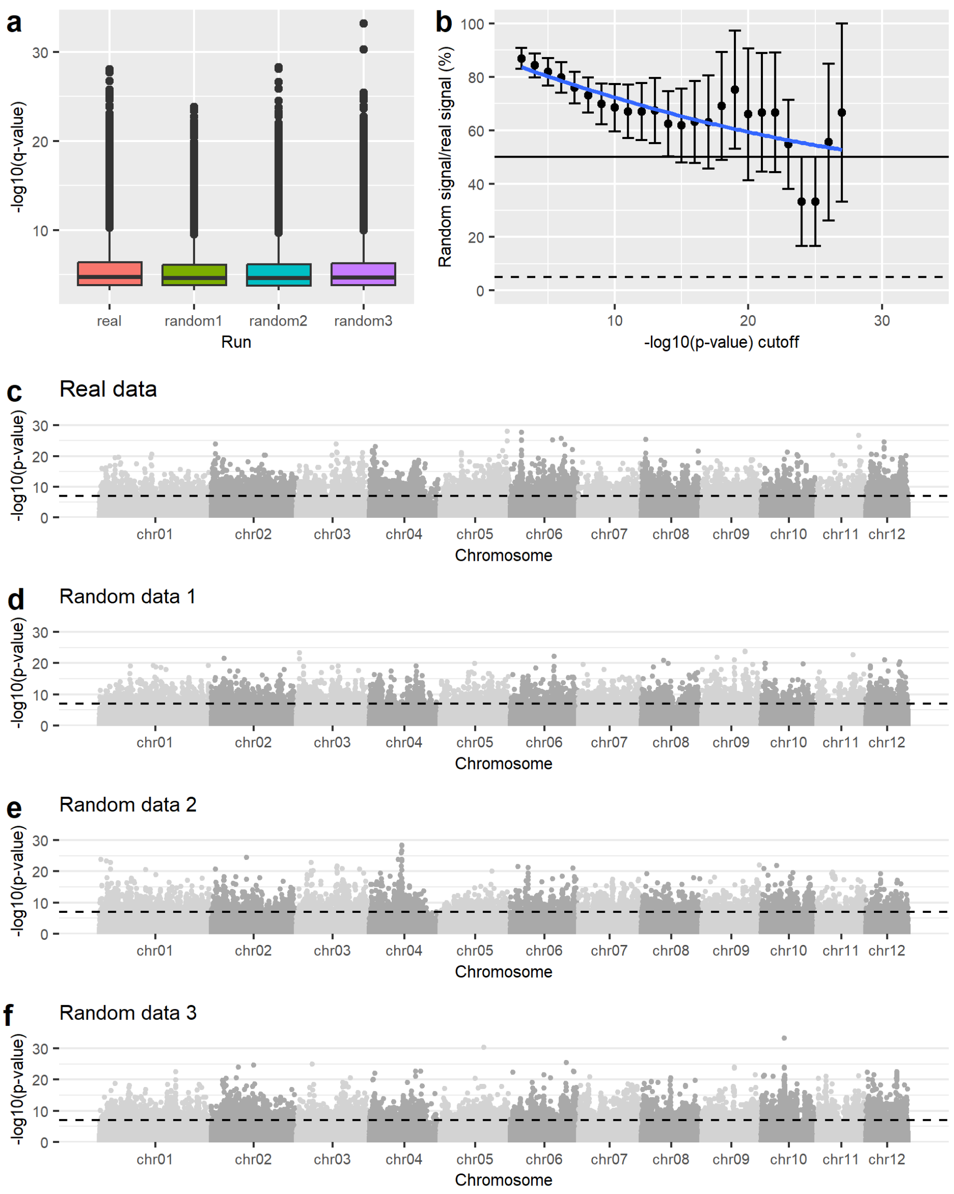


**Supplementary Figure 11**

**GEA analysis with the RDA method shows high level of random signal across *p*-value bins**
(**a**) We ran the RDA analysis[36] using the real data and three random datasets, where the population coordinates and thus the bioclimatic variables were randomly assigned. (**b**) The ratio of random vs. real signal (i.e., the number of associations identified with random data divided by the number of associations identified with the real data) in bins of increasing size, starting with the most significant associations and then adding less significant ones consecutively in -log10(p-value)-steps of one (-log10(p) bins = 27-28, 26-28, 25-28, 24-28 etc.) Blue line shows polynomial regression. (**c**-**f**) Manhattan plots for real (c) and random (e-f) data shows -log10(*p*-values) for 540k independent genome-wide variants in 653 individuals from 98 populations. Chromosomes are distinguished by alternating colors. Dashed horizontal line indicates 5% significance threshold after Bonferroni correction for multiple testing.

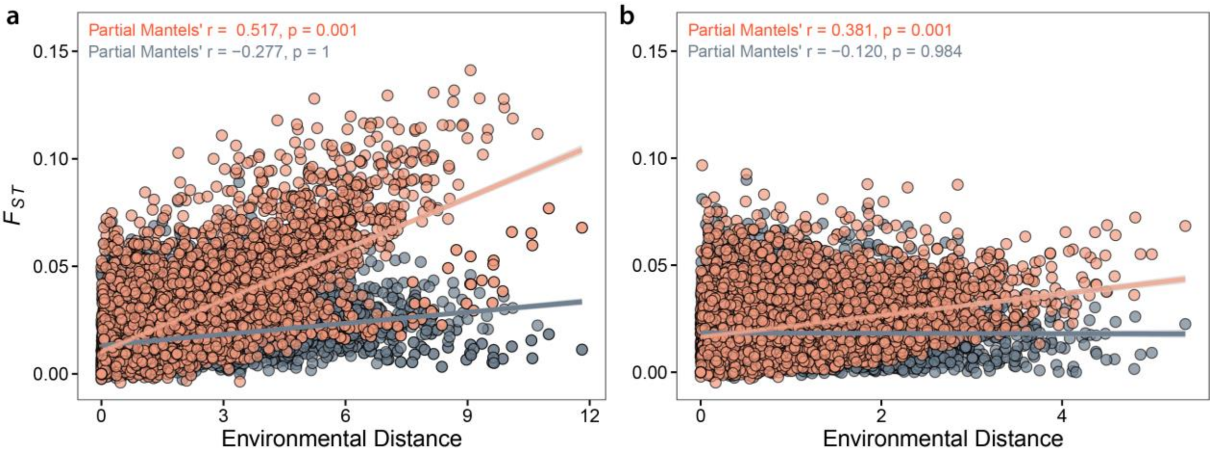


**Supplementary Figure 12**

**Isolation-by-environment (IBE) for associated and non-associated variants using real and random data**

(**a**,**b**) Partial Mantel test assesses the relationship between genetic distance (y-axis) and environmental distance (x-axis) while controlling for the influence of geographical distance. Environmental distance is calculated as the Euclidian distance based on the minimum temperature of the coldest month (bio6) variable. Each dot represents a pairwise comparison among the 98 populations in orange for associated and blue for non-associated variants. The shadow of linear regression denotes the 95% confidence interval of the model for the real (a) and randomized data (b).

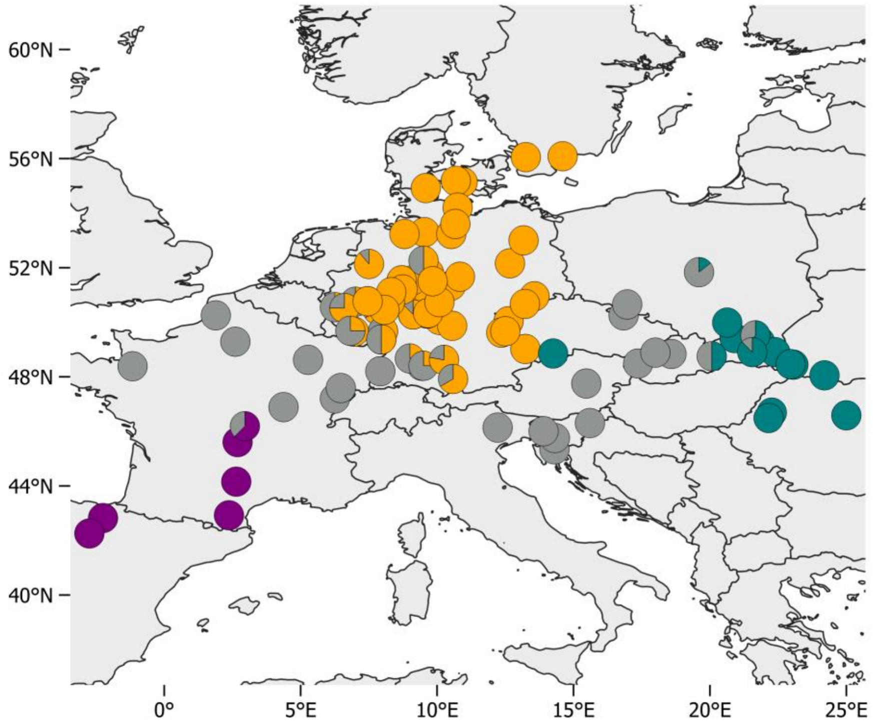


**Supplementary Figure 13**

**Fraction of individuals with more than 70% cluster membership.**

Pie charts represent the genetic composition of 98 populations comprising a total of 653 individuals. Each individual is categorized into one of the three major ancestral genetic clusters (K=3, Supplementary Table 8), determined by a 70% cluster membership probability threshold. Samples predominantly assigned to the western, central, and eastern clusters are shown in purple, orange, and green within the pie chart segments. Individuals failing to meet the 70% threshold are shaded in grey.

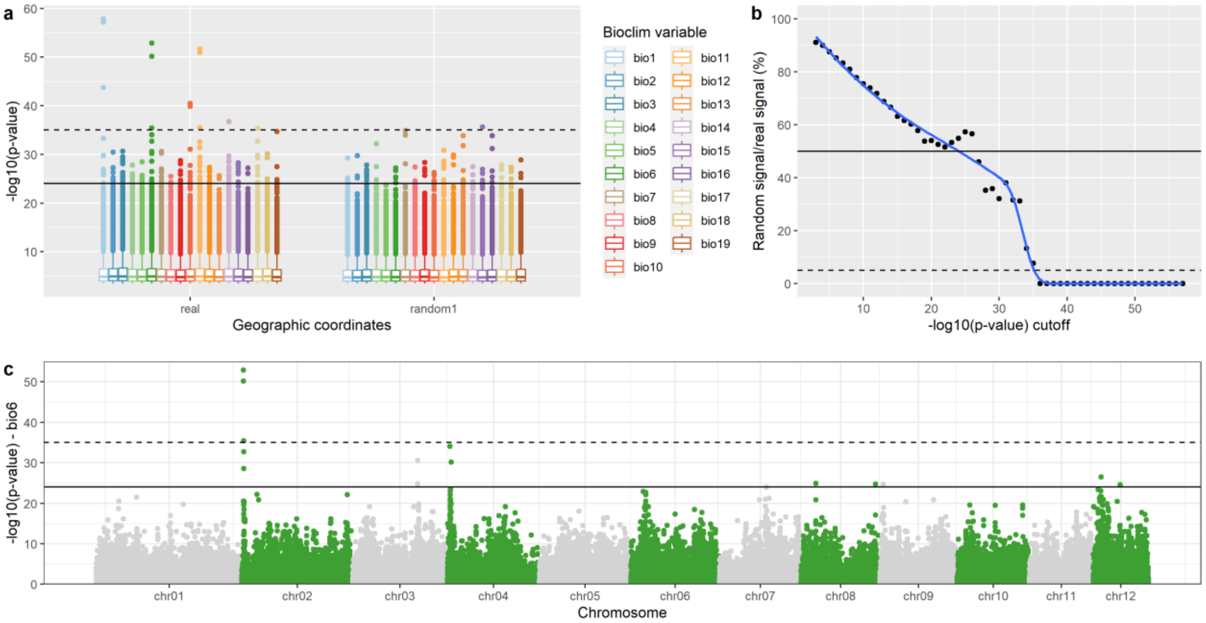


**Supplementary Figure 14**

**LFMM analysis with real and randomized data of local populations revealed the same locus on chromosome 2 as the global analysis (Fig. 3).**

(**a**) We ran a latent factor mixed model (lfmm) analysis[33] using real and randomized population coordinates for populations with at least three individuals with a membership to the central genetic cluster of more than 70% (Supplementary Fig. 12). We performed three independent randomizations. Two variants, especially associated with variation in bio1, bio6 and bio11, stood out. Dashed and solid horizontal lines indicate significance thresholds for which the likelihood of being exceeded with random data compared to the real data is 5% (number of associations with random data/number of associations with real data = 1/20) or 50%, respectively, as determined by comparison of the p-value distributions between real and random data shown in (b). (**b**) Significance thresholds were defined by assessing the ratio of random vs. real signal (i.e., the number of associations identified with random data divided by the number of associations identified with the real data) in bins of increasing size, starting with the most significant associations and then adding less significant ones consecutively in -log10(p-value)-steps of one. Blue line shows a polynomial regression. (**c**) Manhattan plot of the 540k LD-filtered genome-wide variants in 334 individuals from 51 populations (Supplementary Table 8). The significance of the association of each variant along the 12 beech chromosomes with the minimum temperature of the coldest month (bio6) of the real population origins is indicated by the -log10 *p*-values on the y-axis. The dashed and solid horizontal lines indicate significant thresholds as determined in (b).

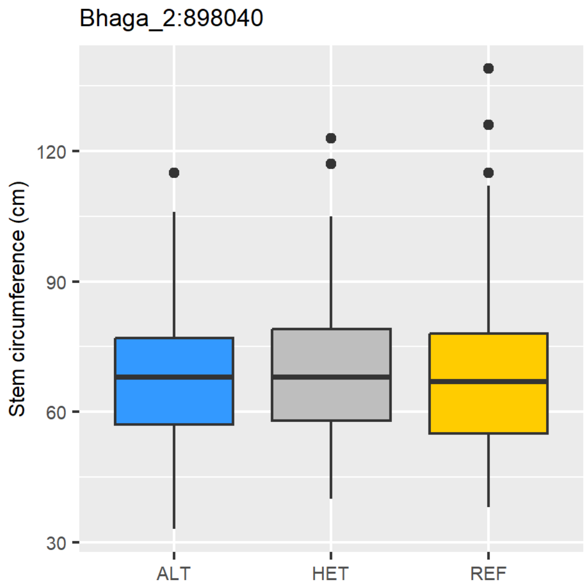


**Supplementary Figure 15**

**Allelic variation for the winter cold locus on chromosome 2 does not affect growth in our common garden**

Boxplots show stem circumference (cm) for the unrelated beech individuals in the common garden ordered by their genotype for the variant Bhage_2:898040 most highly associated with winter cold in an lfmm analysis (ALT: n=157, HET: n=265, REF: n=229). Different genotypes do not differ in stem circumference (one-way ANOVA, $p$=0.51).

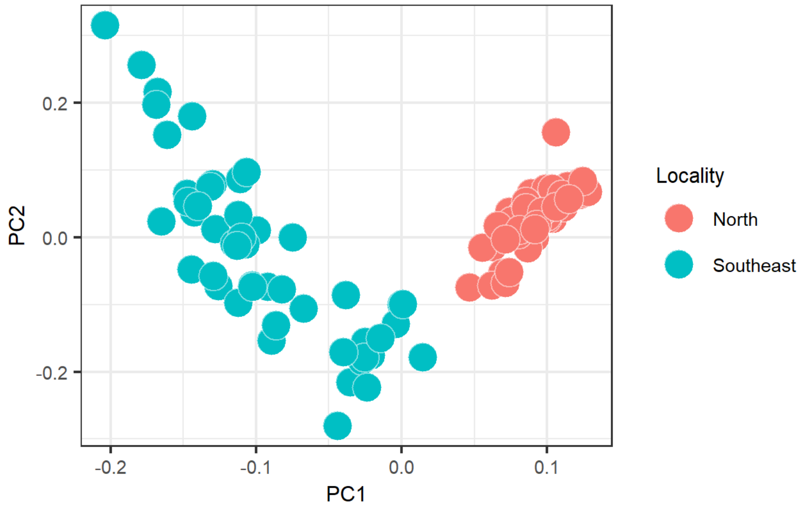


**Supplementary Figure 16**

**Local provenances from the North and the Southeast exhibit consistent genetic differentiation**
Principal component analysis (PCA) using 540k independent genome-wide variants in 98 largely unrelated (less than 2nd degree) individuals from 16 populations, eight from the North and eight from the Southeast (Fig. 4), shows genetic differentiation between localities. Each point represents one of the 98 individuals colored according to their geographic origin.

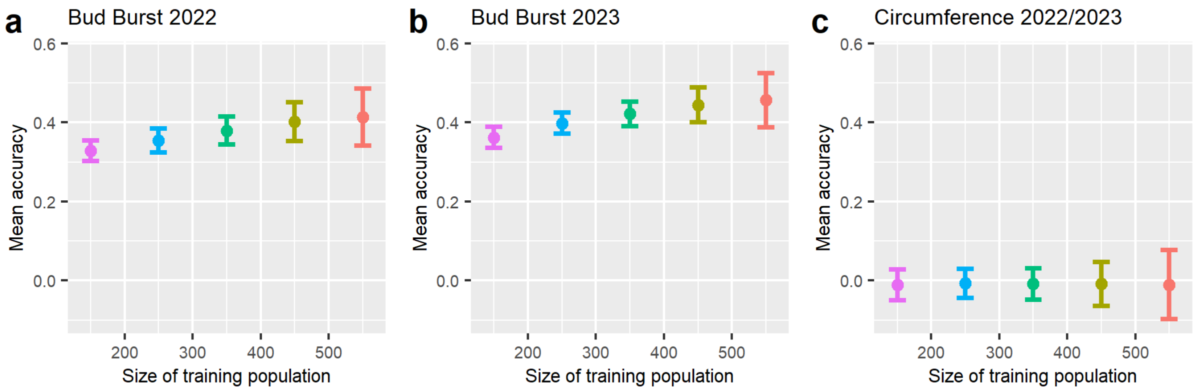


**Supplementary Figure 17**

**Genomic prediction for bud burst in 653 unrelated individuals shows high predictive accuracy.**
(**a**-**c**) Mean genomic prediction accuracies ±SD for 200 cross validations using different sizes of training populations, indicated on the x-axis, are shown for bud burst determined in 2022 (a) and 2023 (b) and stem circumference measured in winter 2022/2023 (c) for the final set of 653 unrelated individuals from 98 populations growing in the common garden in northern Germany. While circumference cannot be predicted in our set of individuals, mean predictive accuracies reach up to 0.45 for bud burst.

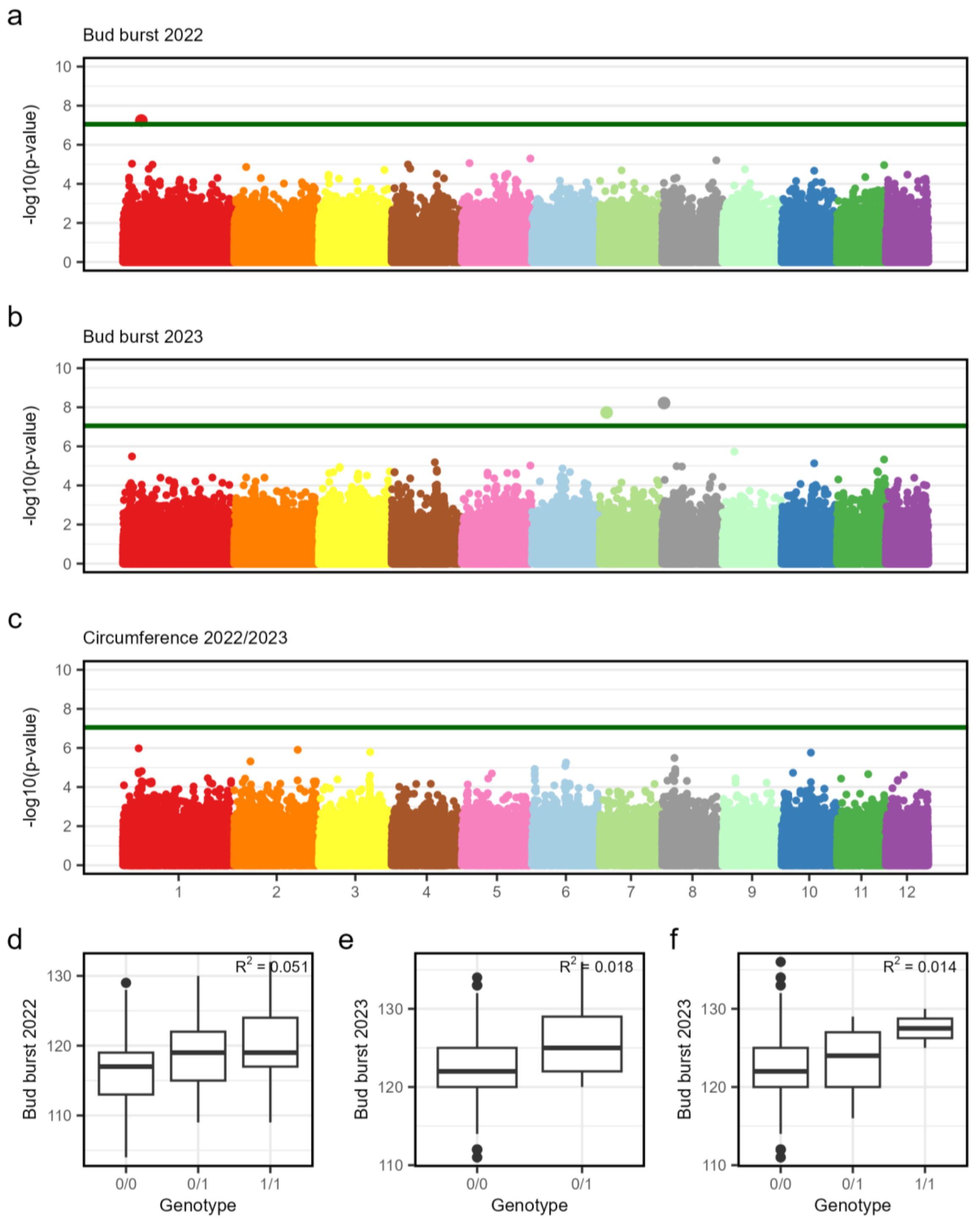


**Supplementary Figure 18**

**Genome-wide association studies (GWAS) reveal polygenicity and missing heritability.**

(**a-c**) We performed GWAS for bud burst assessed for two years, 2022 (a) and 2023 (b), and for growth represented as stem circumference (c) in the final set of 653 unrelated individuals. Green lines on Manhattan plots indicate the Bonferroni-corrected significant threshold and different colors represent different chromosomes. (**d-f**) Boxplots show the relationship between genotype and bud burst for three significant SNPs identified for bud burst in 2022 on chromosome 1 (d), in 2023 on chromosome 7 (e), and in the same year on chromosome 8 (f), with $R^2$ values calculated using one-way ANOVA.

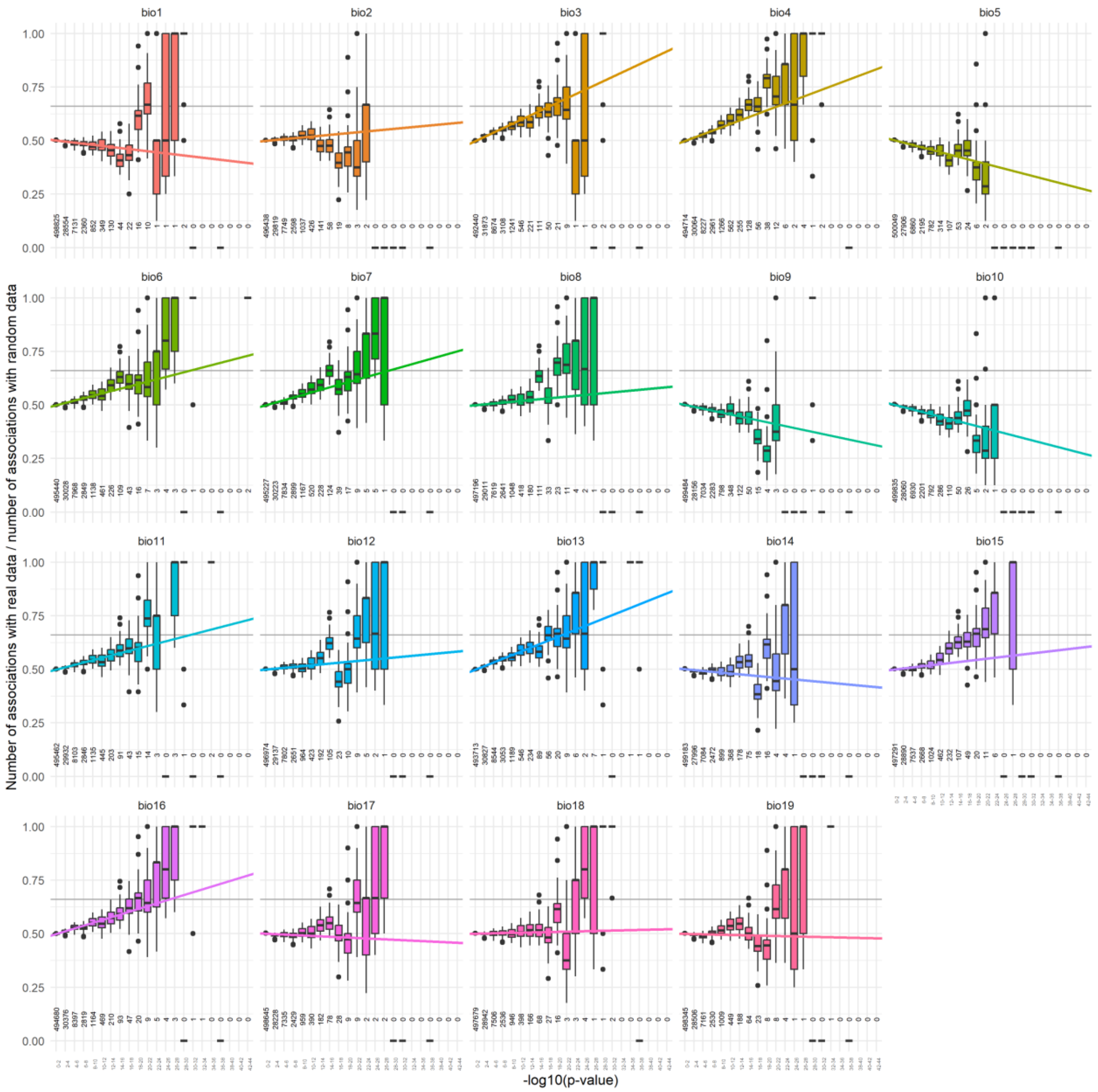


**Supplementary Figure 19**

**Comparison of real vs. random GEAs highlights polygenic local adaptation for some bioclimatic variables**

The fraction of LFMM associations using real and randomized data was analyzed for 22 *p*-value bins (-log10(*p*) of 0-2 to 42-44) for each bioclimatic variable. The number of random associations per p-value bin was determined for three independent data randomizations. The colored line indicates a linear model weighted by the number of associations in each bin, which are given for each bin as vertical numbers underneath the boxes. The horizontal lines mark a 2:1 (0.66) real vs. random fraction, which indicates that the number of associations using the real data is two times the number of associations when using random data. The response to some bioclimatic variables, such as bio3, shows indications of polygenic adaptation with an increasing number of real associations compared to random associations already for *p*-value bins of relatively low significance. These are not clustered into few individual loci but spread across the genome (Supplementary Fig. 8).

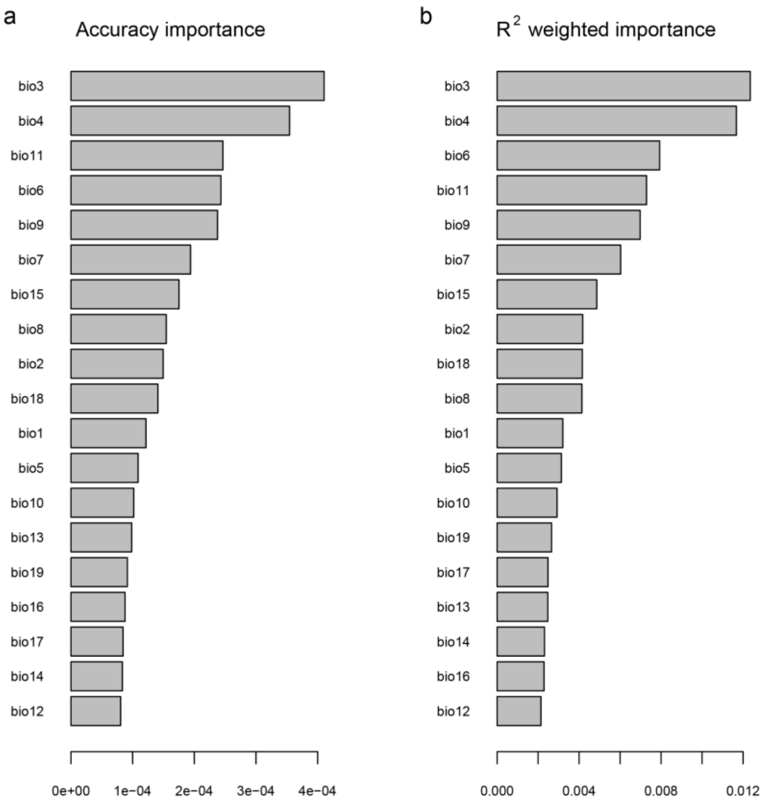


**Supplementary Figure 20**

**Importance ranking of the 19 bioclimatic variables by gradient forests analysis.**

(**a**,**b**) The predictor overall importance as generated by 'gradientForest', for the 19 bioclimatic variables, here ordered according their accuracy importance (a) where importance of each variable is ranked for their prediction accuracy and by mean importance of each variable weighted by SNPs $R^2$ (b).

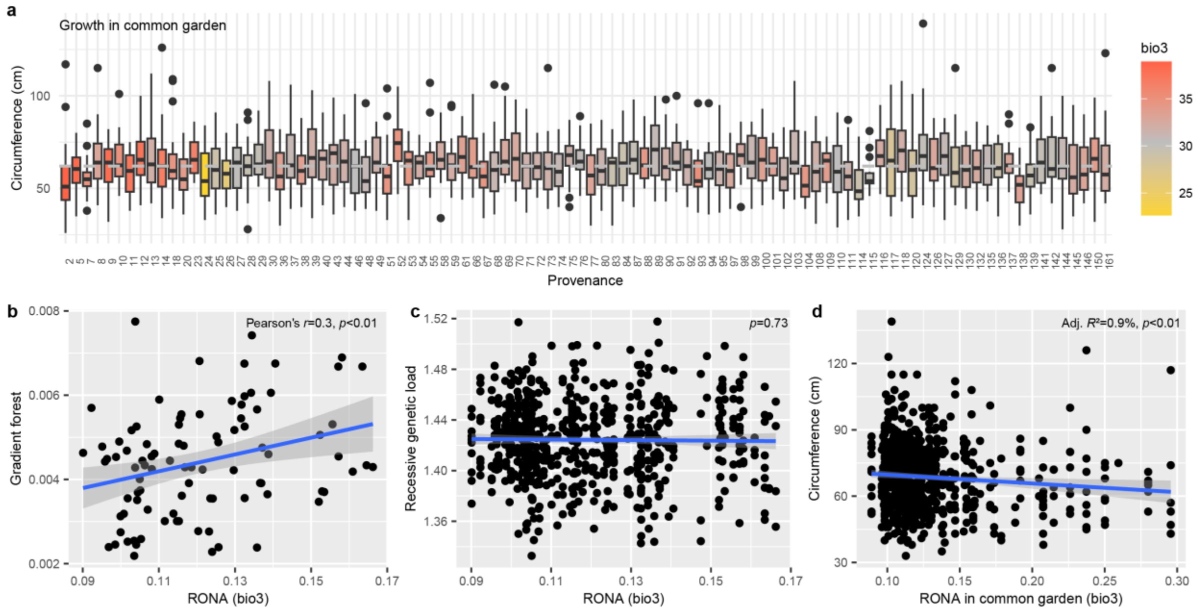


**Supplementary Figure 21**

**Correlations between different genomic offset measures, genetic load and growth in the common garden**

(**a**) Boxplots show stem circumference of the 653 sequenced unrelated trees for the 98 populations. Boxes are colored according to bio3 of the populations' origins. Only a small fraction of the phenotypic variance is non-significantly explained by provenance (one-way ANOVA, adjusted $R^2$=1.8%, $p$=0.21). (**b**,**c**,**d**) Scatterplots show the relationship between RONA and gradient forests offset (b), RONA and recessive genetic load (c), and RONA and growth in our common garden (d). Blue lines and shading indicate linear regression and 95% confidence interval of the model.

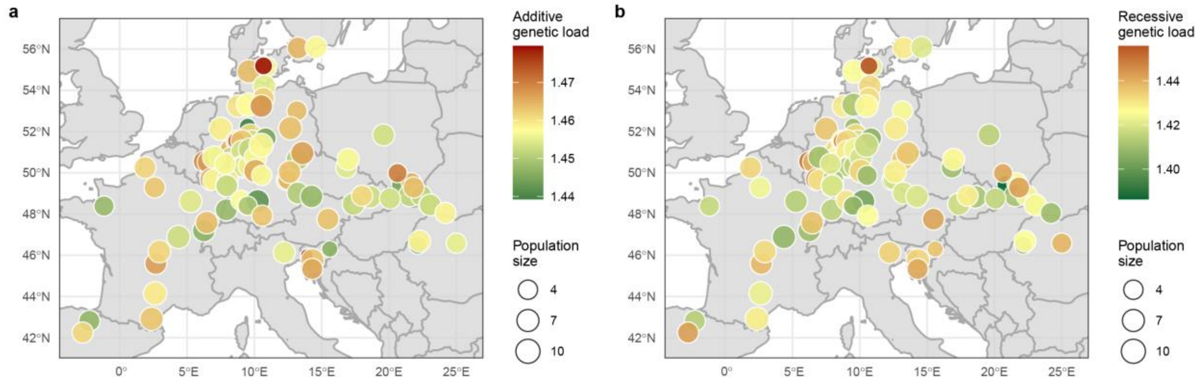


**Supplementary Figure 22**

**Genetic load shows heterogeneous distribution across the landscape.**

(**a, b**) The additive (a) and recessive genetic load (b) is shown for the 98 populations. Colors show average genetic load values, with yellow indicating the median. The size of the circles marks the number of individuals per population (n=4-10). No particular geographical pattern is evident for the two genetic load estimates.

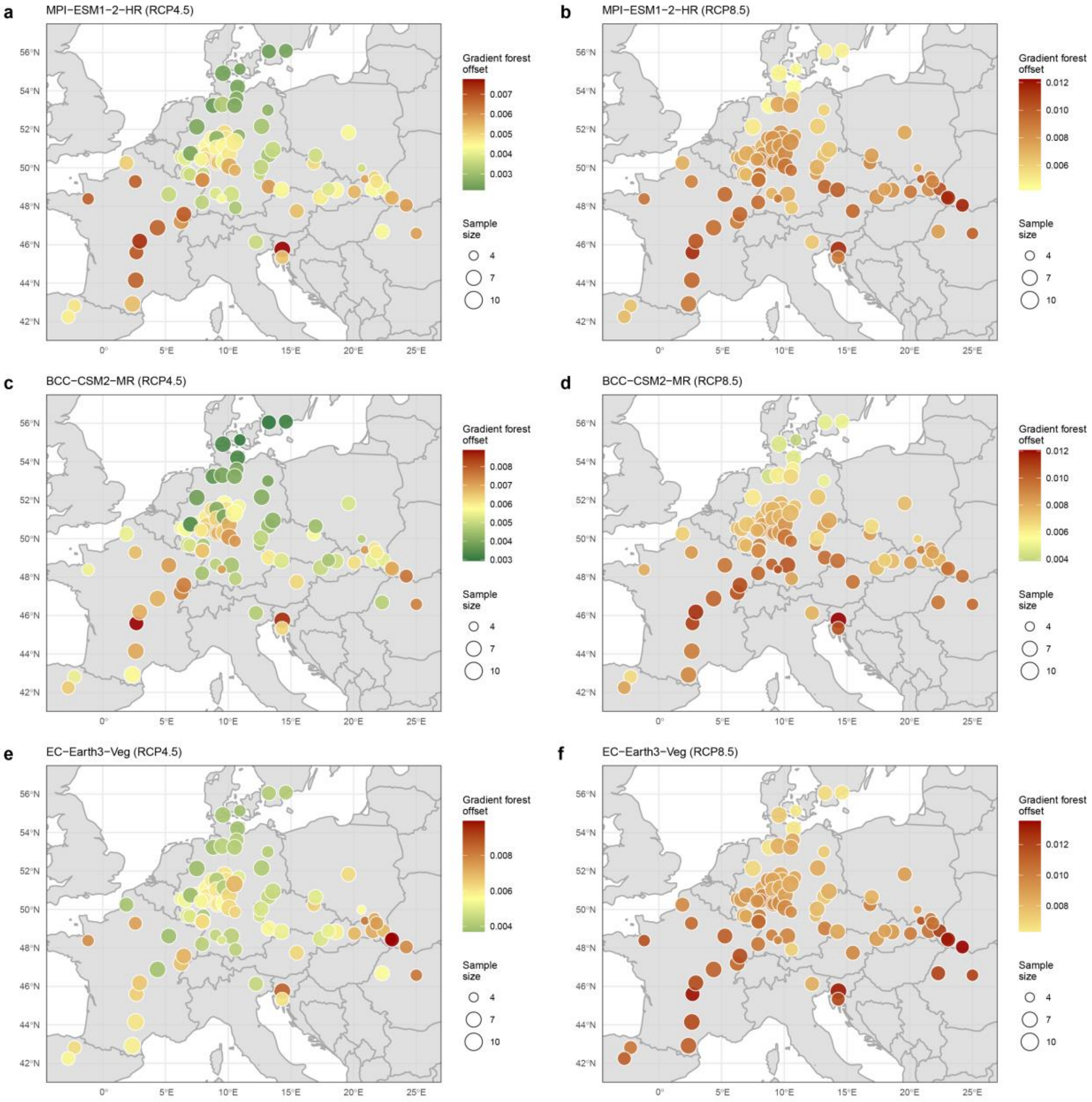


**Supplementary Figure 23**

**Gradient forest offset analyses using different climate change models and scenarios highlight broad- and fine-scale variation**

Gradient forest offset was calculated using 10,000 randomly selected variants and predicted future climate in 2081-2100 compared to near-current (1970-2000) conditions using three different climate change prediction models (‘MPI ESM1 2 HR’ (**a**,**b**) , ‘BCCCSM2-MR’ (**c**,**d**) and ‘EC-Earth3-Veg’ (**e**,**f**)) for two different climate change scenarios (RCP4.5 and RCP8.5). Colors show estimated offset values with yellow indicating the median. For the RCP8.5 scenario, colors are based on the median of the RCP4.5 values. The size of the circles marks the number of individuals per population (n=4-10).

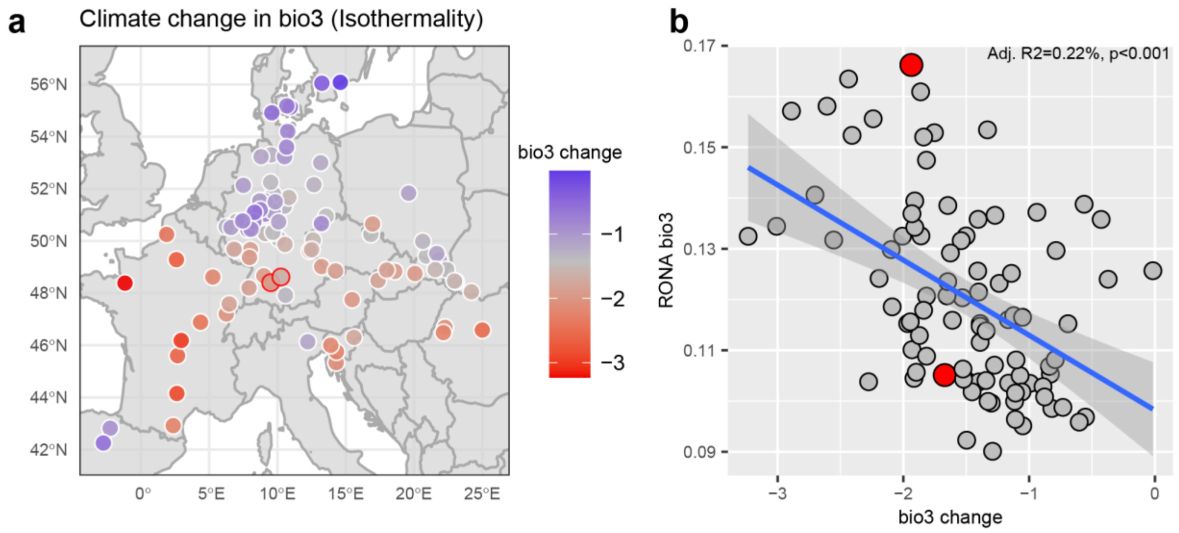


**Supplementary Figure 24**

**Predicted climate change in isothermality (bio3) for the 98 beech populations and comparison between RONA estimates and climate change for bio3.**

(**a**) The map shows the predicted climate change in isothermality (bio3) in 2081-2100 compared to near-current conditions (1970-2000) using an intermediate climate change scenario (RCP4.5, MPI-ESM1-2-HR) for the 98 beech populations analyzed. Circles on the map depict the geographic origins of the populations, colors indicate the predicted climate change with grey representing the median. Two populations from southern Germany, specifically provenance 98 ‘Giengen’ and provenance 99 ‘Ehingen’, representing an example for fine-scale variation in genomic offset, are indicated by red circles (**b**) Comparison of RONA and climate change for bio3 for the 98 populations shown in (a). The two exemplary populations are colored in red. The blue line shows linear model (adjusted $R^2$=0.22, $p$<0.001), the shading indicates the 95% confidence interval of the model.

**Supplementary Table 4**

Partial RDA analyses partition the genetic variance by climate, population structure and geography.

| RDA model | Proportion of total inertia | *P*-value | Proportion of explainable inertia |
|---|---|---|---|
| Full model: F ~ clim. + geog. + struct | 0,36 (*R2*) | <0.001*** | 1.00 |
| Pure climate: F ~ clim. \| (geog. + struct.) | 0,11 (*R2*) | <0.001*** | 0.31 |
| Pure structure: F ~ struct. \| (clim. + geog.) | 0,12 (*R2*) | <0.001*** | 0.33 |
| Pure geography: F ~ geog. \| (clim. + struct.) | 0,02 (*R2*) | <0.01** | 0.06 |
| Confounded climate/structure/geography | 0.11 | | 0.31 |
| Total unexplained | 0.64 | | |
| Total inertia | 1.00 | | |

**Legends for Supplementary Tables 1-3 and 5-9**

Supplementary Table 1: Latitude and longitude for the 100 beech populations planted in the common garden in Schädtbek (trial code: BU1901). Population numbers indicate provenance codes[51].

Supplementary Table 2: Sequencing statistics for the 874 samples for which Illumina data were generated.

Supplementary Table 3: Information on the 653 unrelated trees used for the population structure and genotype-environment association analyses. Block, row and tree indicate the position within the common garden, country the origin of the population.

Supplementary Table 5: Phenotypic measurements for bud burst in 2022, bud burst in 2023 and stem circumference in winter 2022/2023 are given for the 653 unrelated trees in the common garden in Schädtbek, northern Germany (BU1901).

Supplementary Table 6: Diameter at breast height (DBH) measurements for the 16 local provenances in the two common gardens (BU1901 and BU1905).

Supplementary Table 7: Survival measurements for the 16 local provenances in the two common gardens (BU1901 and BU1905).

Supplementary Table 8: Ancestry coefficients as determined by SNMF analysis for the 653 unrelated individuals with K=3.

Supplementary Table 9: Primers used for qRT-PCR of the Callose synthase 1 gene (Bhaga_2.g94).
